# Contrasting defense strategies of oligotrophs and copiotrophs revealed by single-cell-resolved virus–host pairing of freshwater bacteria

**DOI:** 10.1101/2024.07.24.604879

**Authors:** Yusuke Okazaki, Yohei Nishikawa, Ryota Wagatsuma, Haruko Takeyama, Shin-ichi Nakano

## Abstract

The ecological importance of virus–host interactions is unclear due to the limited ability of metagenomics to resolve virus–host pairs and the infection state of individual cells. We addressed these problems using single-cell genomics combined with published metagenomic data on lake bacterioplankton. We obtained 862 medium- to high-quality single-cell amplified genomes (SAGs) from two water layers and two seasons in Lake Biwa, Japan. We assembled 176 viral (dsDNA phage) contigs in the SAGs, and identified novel virus–host pairs including the discovery of viruses infecting CL500-11, the dominant bacterioplankton lineage in deep freshwater lakes worldwide. A virus was detected in 133 (15.4%) SAGs through read mapping analysis. The viral detection rate showed little variation among samples (12.1–18.1%) but significant variation in host taxonomy (4.2–65.3%), with copiotrophs showing higher values than oligotrophs. The high infection rates of copiotrophs were achieved by collective infection by diverse viruses, suggesting weak density-dependent virus–host selections, presumably because of their non-persistent interactions with viruses due to their fluctuating abundance. In contrast, the low infection rates of oligotrophs supported the idea that their co-dominance with viruses is achieved by genomic microdiversification that diversifies the virus–host specificity, sustained by their large population size and persistent density-dependent fluctuating selection. Overall, we demonstrated that virus–host interactions are highly diverse within and between host lineages, which was overlooked by metagenomics analysis, as exemplified by the CL500-11 virus, which showed extremely high read coverages in cellular and virion metagenomes, but infected < 1% of host cells.

**Significance statement:** Virus–host interactions are among the most significant driving forces of microbial biogeochemical cycles and genomic diversification. Unlike experimental conditions, bacterial cells in the natural environment are not uniformly infected by a single virus, but interact with diverse viruses under heterogeneous eco-physiological and genetic conditions. The specificity and heterogeneity of infection are the keys to understanding complex virus–host interactions and the mechanisms behind their co-existence. However, these interactions remain unclear due to the limitations of conventional metagenomic approaches. We addressed this issue by detecting viral signals from single-cell-amplified genomes of lake bacterial communities. The results revealed novel virus–host pairs and their infection rates, suggesting that viral defense strategies differ among host lineages, reflecting their ecological characteristics.

## Introduction

Metagenomics has revealed the enormous diversity of microorganisms and their viruses in the environment, leading researchers to focus on their ecological importance. Viruses are major mortality factors of bacteria (1) and primary drivers of genome diversification resulting from a complex virus– host arms race (2). Although metagenomics allows the recovery of thousands of uncultured microbial and viral genomes from the environment, virus–host interactions remain unclear due to two technical limitations. The first limitation is the difficulty of host prediction. Much effort has focused on linking virus and host using metagenomic information, for example, through sequence matching, nucleotide frequency, similarity to isolated viruses, lineage-specific marker genes, and co-occurrence with potential hosts (3, 4). However, most metagenome-derived viral genomes lack such information to confidently predict the host. The second limitation is the difficulty in resolving heterogeneous viral infection status among individual host cells. Unlike experimental conditions, cells of a natural host population are not uniformly infected by viruses due to their physiological and genetic heterogeneity. Indeed, host genomic microdiversity is crucial to understanding complex virus–host interactions (2, 5). However, metagenomics overlooks such intra-population heterogeneity because it targets populations of cells and virions.

To overcome the limitations of metagenomics, we conducted single-cell genomic analyses of environmental bacterial assemblages. We targeted bacterioplankton and their viral community in Lake Biwa, Japan, where metagenomic studies have reconstructed > 500 bacterial and > 7000 viral genomes and characterized their spatiotemporal distributions (6–9). Although these studies identified quantitatively significant bacteria and viruses in the lake, the limited resolution of metagenomics hindered the prediction of virus–host pairs and the elucidation of their interactions. For example, Actinobacteria and their viruses are the most abundant and diverse members of the microbial ecosystem in Lake Biwa and in freshwater lakes globally (6, 10, 11). However, due to methodological limitations, little is known of their virus–host specificity and infection rate in this ecosystem, although they are the keys to understanding the mechanisms behind virus–host co-dominance and their sustainable diversity.

We investigated the interaction between bacteria and their virus (double-stranded DNA phages) at single-cell resolution by identifying viral signals in individual single-cell amplified genomes (SAGs). This approach has been used to identify novel virus–host pairs in marine (12–14) and hot spring (15, 16) systems but not in freshwater lakes. Based on published metagenomic data (7, 8) and high-quality SAGs generated from multiple seasons and depths using state-of-the-art gel bead-based SAG (SAG-gel) technology (17), we explored virus–host interactions in the lake at single-cell-resolution. The results identified many novel virus–host pairs, including those involving the most quantitatively significant freshwater bacterioplankton lineages, which were previously overlooked in metagenomic studies. We demonstrated the cell-to-cell heterogeneity of viral infection status and compared viral infection rates among samples and host lineages. Specifically, our results highlighted contrasting defense strategies between oligotrophs and copiotrophs.

## MATERIALS AND METHODS

### Sample collection and single-cell genome sequencing

Microbial samples were collected near the deepest point of Lake Biwa, Japan (water depth, ∼101 m) (35°20.10N 136°06.16E) in summer (June 29, 2022) and winter (February 1, 2023). Vertical profiles of temperature and dissolved oxygen concentration (DO) were analyzed *in situ* using a conductivity, temperature, and depth (CTD) probe (Rinko ASTD102; JFE Advantech, Tokyo, Japan). In summer, the water was thermally stratified, with a surface water temperature of 26°C. In winter, the water column was almost holomictic, with an oxycline at ∼85-m depth, where DO decreased from 10.1 to 3.9 mg L^−1^ (Fig. S1). In both seasons, we collected two samples representing the epilimnion and hypolimnion (5 and 80 m in summer and 5 and 95 m in winter). For each sample, 1 L of lake water was collected and kept under cool, dark conditions and processed within 24 h. Concentration, gel bead encapsulation, DNA extraction, whole-genome amplification, fluorescent staining, sorting, and sequencing of individual cells were performed following a published SAG-gel protocol (17). For each sample, we sorted 192 and 768 gel beads in summer and winter, respectively, for a total of 1920 gel beads. For each gel bead, 0.6–546.6 Mb of reads (median = 46.3 Mb) were generated by Illumina NextSeq 2000 paired-end sequencing (2 × 150 bp). For each assembly, we labeled the samples “LB2206” for summer and “LB2301” for winter, and appended an E or H for epilimnion or hypolimnion samples, respectively, followed by a numerical identifier.

### Generation and taxonomic annotation of SAGs

The raw reads from each gel bead were preprocessed by bbduk v38.96, using the parameters ktrim=r ref=adapters k=23 mink=11 hdist=1 tpe tbo qtrim=r trimq=10 minlength=40 maxns=1 minavgquality=15 (https://sourceforge.net/projects/bbmap/). Assembly was conducted using SPAdes v3.15.2, using the parameters --sc --careful --disable-rr (18). Using SeqKit v2.4.0 (19), we removed < 1-kb contigs and calculated the N50 of each assembly. The quality and taxonomy of the genomes were evaluated by CheckM v1.2.2 (20) and GTDB-Tk v2.2.6 (reference data version = r207) (21, 22), respectively. In the analysis, we replaced the GTDB family name “UBA11657” with “CL500-11”, which is a familiar name for this uncultured bacterioplankton lineage (23, 24). Based on the CheckM results, we defined a quality score (QS) for each assembly as “Completeness – 5 × Contamination”, which is a metric commonly used to evaluate the quality of metagenome-assembled genomes (MAGs) (22, 25). Assemblies with QS > 50 and QS = 30–50 were defined as high- and medium-quality SAGs, respectively. In total, 862 medium- or high-quality SAGs were obtained and used in downstream analyses. The circularity of an assembly was tested using ccfind v1.4.5 (26). A pangenome graph of SAGs was generated using SuperPang v1.1.0 (27) using the default parameters, and visualized by Bandage v0.9.0 (28).

### Identification of viral contigs in a SAG

We used geNomad v1.5.1 (29), VIBRANT v1.2.1 (30), and VirSorter2 v2.2.3 (31) to identify dsDNA phage contigs in each SAG. In geNomad, we used the “end-to-end” pipeline with the default parameters, and all contigs annotated as Caudoviricetes were flagged. In VIBRANT, we used the “VIBRANT_run.py” script with the default parameters, and contigs reported as phages were flagged. In VirSorter2, we used the “virsorter run” pipeline with default parameters, and contigs with a dsDNAphage score > 0.7 were flagged. Next, contigs flagged by two or three tools were selected and processed for quality control using CheckV v1.0.1 (32). If CheckV reported “no viral genes detected” in a contig, the contig was discarded. If CheckV reported a contig as proviral, the viral segment was extracted from cellular regions and used in downstream analyses. Gene prediction, functional annotation, and alignment of the viral contigs were performed and visualized using the ViPTree v4.0 (33) and DiGAlign v2.0 (34) webservers. Further high-sensitivity searches for viral hallmark genes were conducted using SDsearch v0.2.0 (26) with the default parameters against the pfamA hhsuite v35.0 database (35).

### Generation of a reference database from published metagenomic data

We used metagenomic datasets collected by two published studies from Lake Biwa to comprehend the viral genomic diversity in the lake. The first (Lake Biwa viral contigs, LBVC) was reconstructed from cellular (0.2–5 μm size fraction) and virion (< 0.2 μm) metagenomes collected monthly from June 2016 to February 2017 from the epilimnion and hypolimnion (6). The second (Lake Biwa viruses, LBV) was reconstructed from virion metagenomes collected monthly from September 2018 to December 2019 from the epilimnion and hypolimnion (8). Contigs > 10 kb were selected from both datasets and quality-controlled using CheckV, as described above. Finally, the metagenome-derived and SAG-derived viral contigs were pooled and dereplicated using dRep v3.4.2 (36) at the 95% average nucleotide identity (ANI) threshold (-sa 0.95 --ignoreGenomeQuality --S_algorithm ANIn -l 1000). The resulting 7,978 non-redundant viral contigs were used as a reference in read-mapping analyses. Phylogenetic relatedness among the reference contigs was tested by tBLASTx-based homology searching using ViPTreeGen v1.1.3 (33).

### Read mapping analyses

Read mapping and coverage calculation were performed using CoverM v0.6.1 (https://github.com/wwood/CoverM). Raw reads of each SAG were mapped to the corresponding SAG to determine the assembly coverage of the contigs. The raw reads were also mapped to the 7,978 reference viral contigs to test their presence in each SAG. In this study, a virus was defined as “detected” when > 50% of the viral contig was covered by the mapped reads. We mapped reads of published metagenomes to the viral contigs to evaluate their spatiotemporal distribution in the lake. We used two datasets: virion (< 0.2 μm) metagenomes collected monthly from December 2018 to November 2019 from the epilimnion and hypolimnion (8), and cellular (0.2–5 μm) metagenomes collected monthly from May 2018 to April 2019 (7). For metagenomic read mapping, the CoverM option “--min-read-percent-identity 92” was used. Metagenomic read coverages were compared among samples using reads downsampled to one million pairs to normalize the sequencing depths.

## RESULTS

### SAG quality and identification of viral contigs

Among the 1920 sequenced gel beads, 1657 resulted in an assembly with a > 1-kb contig. The assembly N50 ranged from 1.0 to 1246.7 kb, and QS ranged from < 0 to 100 with a median of 31.75 (Fig. 1A, Table S1). In total, 528 were high-quality SAGs (QS > 50), and 334 were medium-quality SAGs (QS = 30–50) (Fig. 1B); these were used in downstream analyses. Notably, a 1.15-Mb SAG (LB2301E_00210, genus *Fonsibacter*) was circularity assembled with a QS of 100. The proportions of medium- and high-quality SAGs varied among the samples from 22.3% in the winter hypolimnion to 73.4% in the summer epilimnion (Fig. 1B). Regarding phylogenetic affiliation, eight families (CL500-11, Pelagibacteraceae, Nanopelagicaceae, Ilumatobacteraceae, Burkholderiaceae, Beijerinckiaceae, Chitinophagaceae, and Methylophilaceae) accounted for > 80% of all SAGs (Fig. 1C). The summer epilimnion was dominated by Nanopelagicaceae and Pelagibacteraceae, whereas CL500-11 predominated in the other samples.

**Fig. 1.**
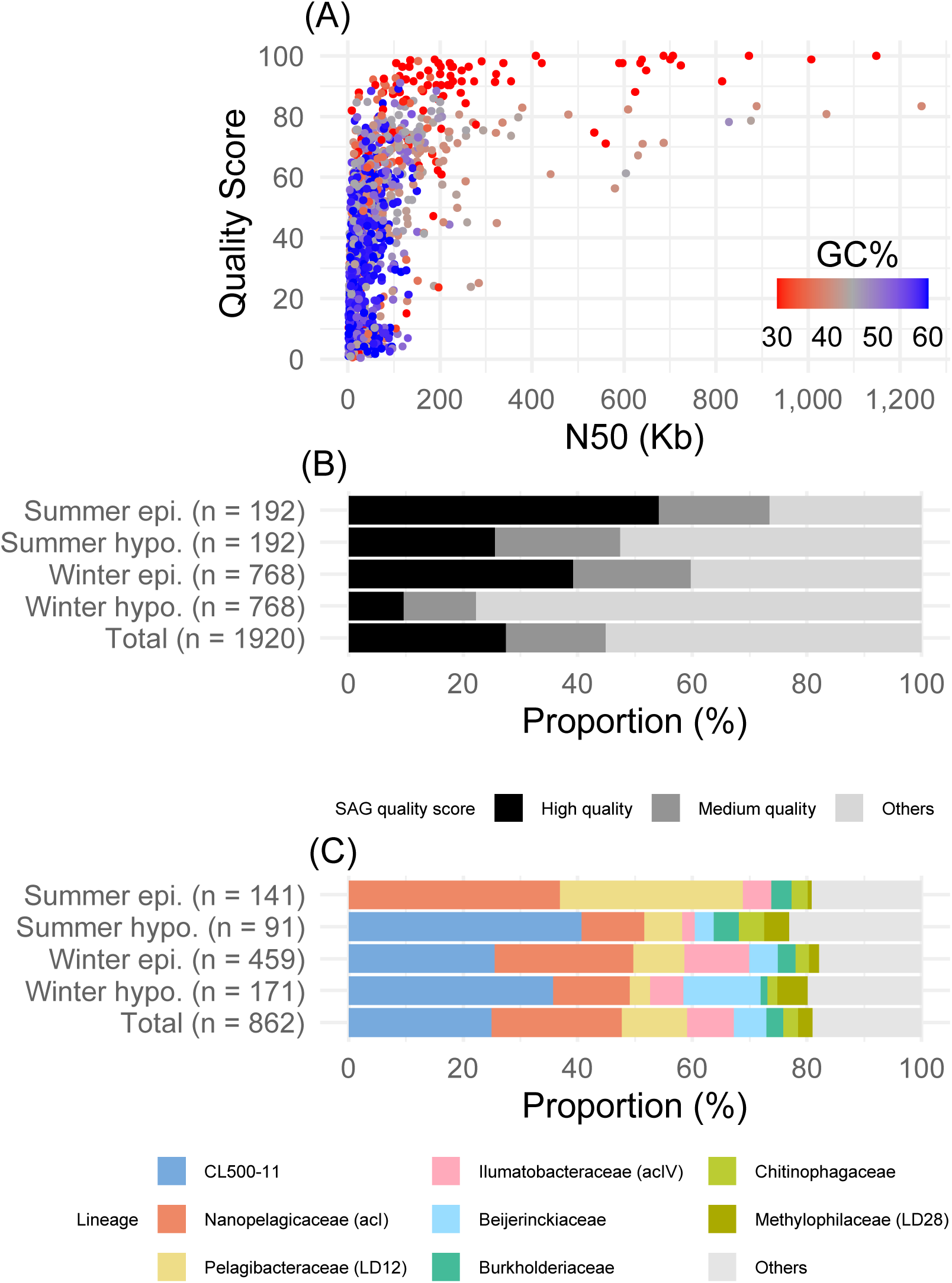
Statistics of single-cell amplified genomes (SAGs). (A) N50 and quality score (QS) distribution of individual SAGs. Colors indicate GC percentages of SAGs. SAGs with QS < 0 are not shown. (B) Quality distribution of SAGs in each sample and in total. High and medium quality are defined by QS > 50 and QS = 30–50, respectively. (C) Phylogenetic composition (family level) of medium- and high-quality SAGs in each sample and in total.

In total, 176 viral contigs were found in 85 SAGs. CheckV reported that 34 of these were proviruses integrated into host genomes. After the excision of the viral region for a provirus, the viral contigs were 1030–118,892 bp in length (Fig. 2A, Table S2). CheckV completeness scores ranged from 0 to 100, with 98 (55.7%) viral contigs showing < 10 and 24 viral contigs (13.6%) showing > 50. Ten viral contigs were reported as complete (completeness score = 100); five of these were identified as proviruses, and the other five as circular contigs (Table S2).

**Fig. 2.**
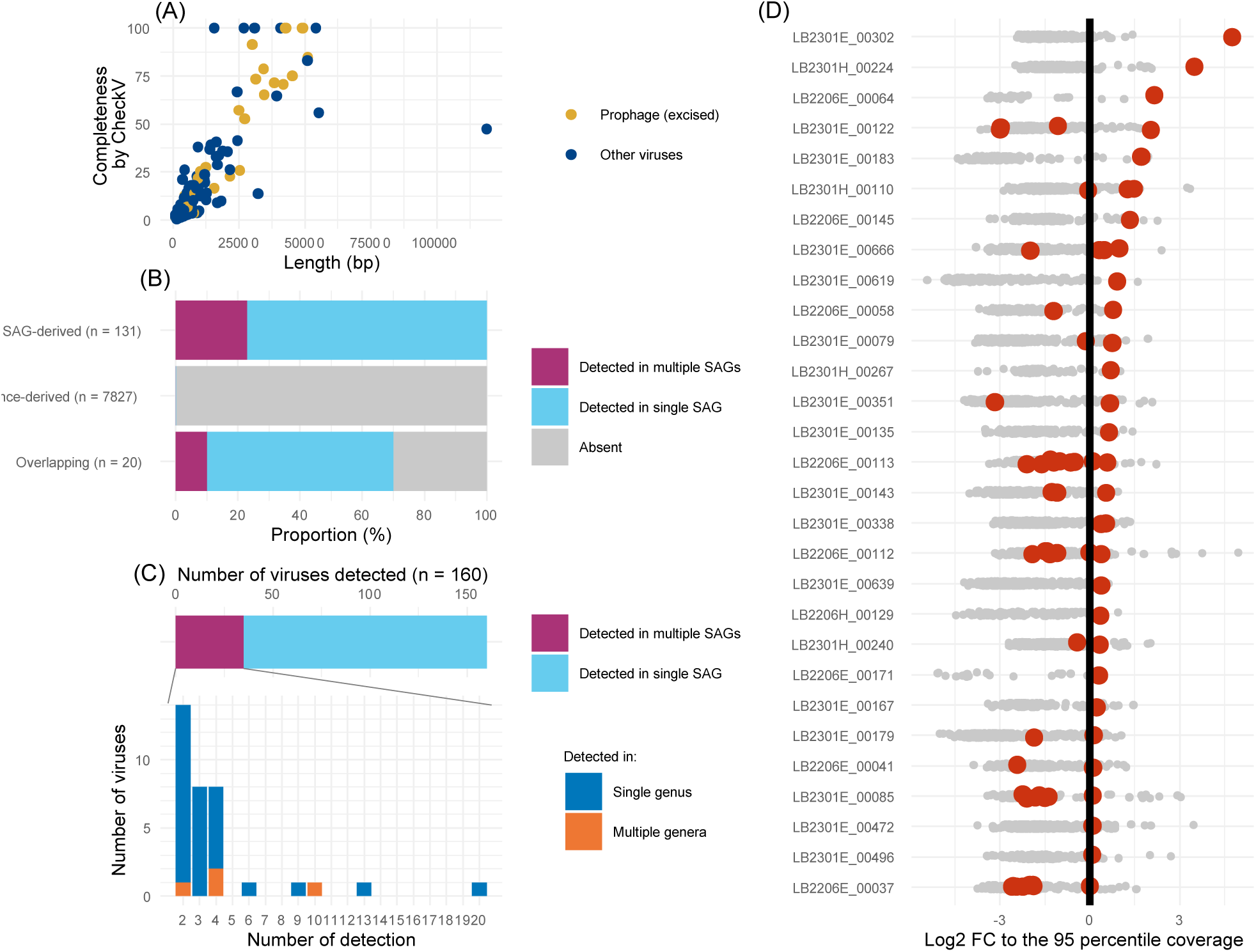
Statistics of detected viruses. (A) Length and completeness distribution of viral contigs assembled in the SAGs. Color indicates whether the contig was detected as prophage (integrated into a host chromosome). Prophage length is defined as the length of the excised viral region. (B) Proportion of viruses detected in SAGs for each clustering category. Different colors indicate detection in single or multiple SAGs. SAG- and reference-derived mean members dereplicated exclusively from viral contigs originating from SAGs and metagenomic reference sequences, respectively, where those derived from both are classified as overlapping. (C) Distribution of the number of viruses detected in different numbers of SAGs. Top panel, proportions of SAGs in which single and multiple viruses were detected. The total number (n = 160) corresponds to the total detected viruses in Fig. 2B. Bottom panel, distribution of multiple detections; colors indicate whether a virus was consistently detected in the same genus. (D) Distribution of contig assembly coverages in each SAG. Coverage was normalized by log2 fold change to the 95^th^ percentile coverage (vertical solid line) for each SAG. Plots indicate individual contigs; viral contigs are indicated in red. Only SAGs with a viral contig with > 95^th^ percentile coverage are shown, sorted by normalized coverage of the viral contig.

### Viral detection for each SAG through read mapping

The 176 SAG-derived viral contigs were pooled with metagenome-derived contigs and dereplicated at the 95% ANI threshold. The resulting 7,978 non-redundant contigs comprised 131 and 7,827 contigs representing clusters generated exclusively from SAG-derived and metagenome-derived contigs, respectively. The other 20 contigs represented clusters including both members. Mapping of SAG raw reads to the viral contigs demonstrated that the 131 SAG-derived viral contigs were detected in at least one SAG, and 30 were detected in multiple SAGs (Fig. 2B). Among the 7,827 reference-derived viral genomes, 15 were detected, and three were detected in multiple SAGs. Among the other 20 viruses in the overlapping category, 14 were detected, and two were detected in multiple SAGs.

Among the detected viruses, we screened for those with a higher read coverage compared to other contigs in the SAG. We operationally set the 95^th^ percentile of contig coverages in a SAG as the threshold to define high-coverage contigs. We found 29 SAGs with a high-coverage viral contig and inspected their contig coverage distribution. Overall, the coverages were distributed continuously and broadly, with no apparent gap (Fig. 2D). Exceptionally, we observed an extremely high-coverage viral contig in each of two CL500-11 SAGs, LB2301E_00302 and LB2301H_00224. Their read coverages (555 and 324, respectively) were far higher than the second-highest coverage (55 and 120, respectively) and the 95^th^ percentile coverage (20.6 and 28.8, respectively) observed in each SAG.

Among the 160 viral contigs detected by read mapping (Fig. 2B), 125 were detected in a single SAG, and 35 were detected in multiple SAGs (Fig. 2C). Notably, one virus contig (LB2301E_00219_ctg_540) was detected in 20 SAGs of the same genus of CL500-11. Most other viruses detected in multiple SAGs were also consistently detected in the same genus (Fig. 2C).

Viral detection rates among the SAGs in each sample ranged from 12.1% (summer hypolimnion) to 18.1% (winter hypolimnion) (Fig. 3A), and the values in each family ranged from 4.2% (Ilumatobacteraceae) to 65.3% (Beijerinckiaceae) (Fig. 3B). We observed heterogeneous virus– host correspondence within each family (Fig. 4). In the Beijerinckiaceae, for instance, 35 viral contigs were detected from 32 SAGs, and the most widespread virus was detected in less than half of the cells (13 SAGs) (Fig. 4A).

**Fig. 3.**
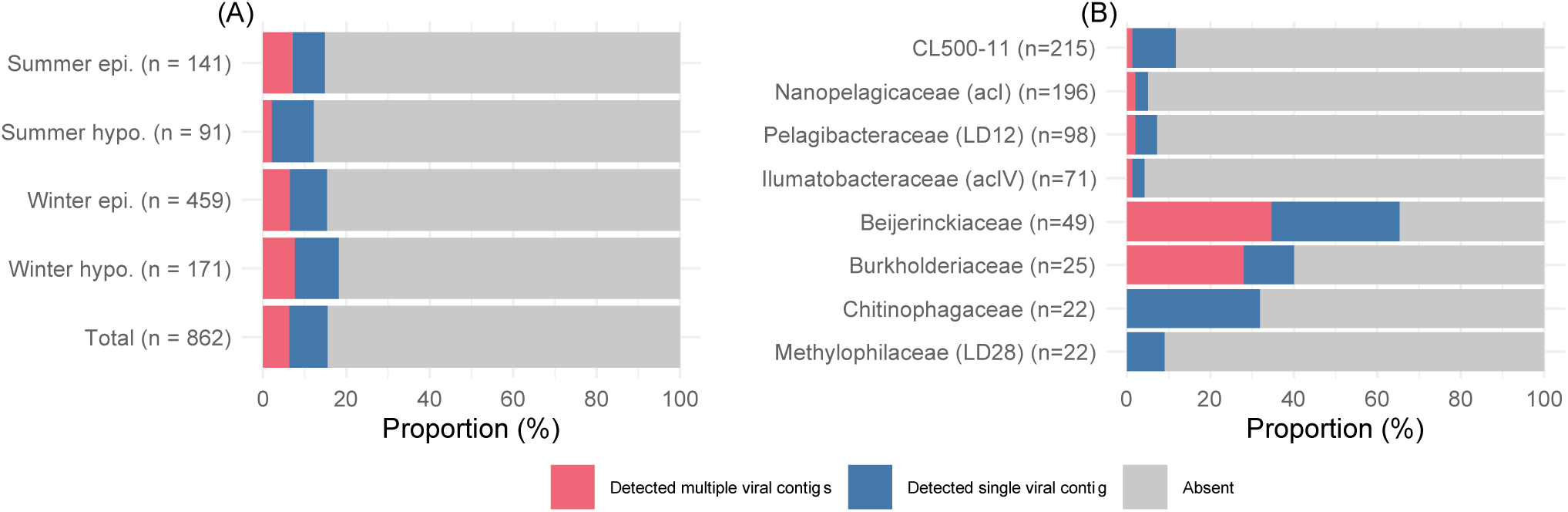
Proportions of SAGs with viral detection. Different colors indicate the detection of single or multiple viral contigs. (A) Proportion in each sample and in total. (B) Proportion in each family.

**Fig. 4.**
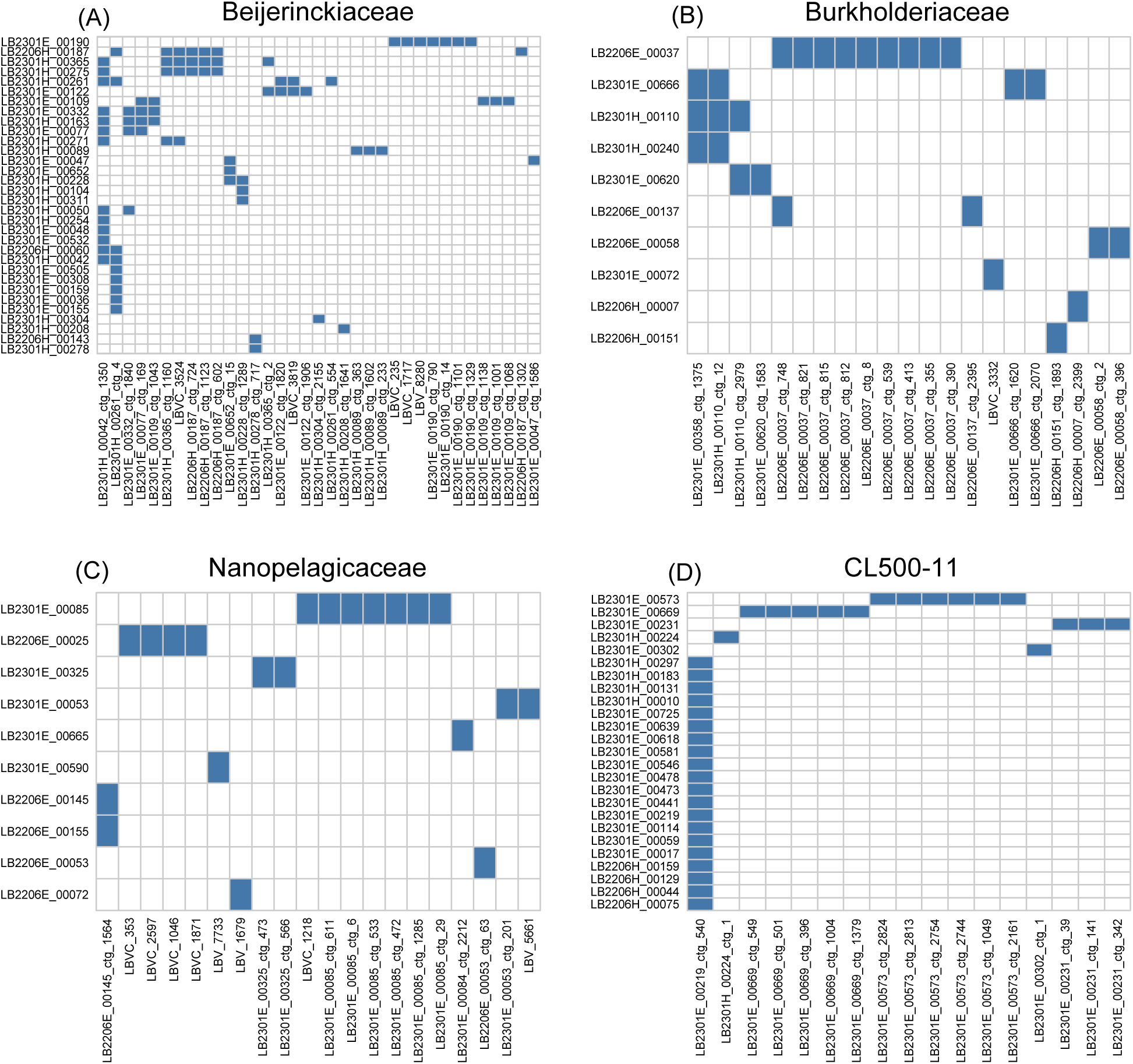
Combinations of SAGs and their detected viral contigs for members of the (A) Beijerinckiaceae, (B) Burkholderiaceae, (C) Nanopelagicaceae, and (D) CL500-11. Horizontal and vertical axes indicate individual viral contigs and SAGs, respectively. Pairs of SAGs and detected viral contigs are indicated by colored boxes.

In 55 SAGs, multiple viral contigs were detected (Fig. 3A), implying co-infection by multiple viruses. We manually inspected the lengths and redundancies of viral marker genes (capsid, terminase, and portal proteins) of the viral contigs detected in the same SAG to determine if they originated from multiple viral genomes or fragments of the same viral genome (Table S3). As a result, one SAG (LB2301E_00122) likely detected contigs of different viral genomes, where marker gene redundancy was observed among multiple >10-kb contigs sharing no homology (Fig. S2). For the other 54 SAGs, the situation was unclear because of fragmented assemblies lacking marker genes (Table S3), except for several cases where the viral contigs originated from the same viral genome that eluded de-replication because the aligned fraction size was lower than the threshold (10%) set in dRep (Fig. S3).

The viral detection rate within each family varied among samples (Fig. S4), but this variation was not statistically supported due to marked differences in the number of SAGs among samples. Although Pelagibacteraceae, Beijerinckiaceae, and CL500-11 harbored one or two species, other dominant families harbored five or more species, with a maximum of 15 species within Nanopelagicaceae (Fig. S5).

### Profiling of the viral distribution in the lake by metagenomic read mapping

Mapping reads of published metagenomes in the lake revealed that among the 131 dereplicated viral contigs originating exclusively from the SAGs, 73% and 24% were detected (showed > 50% coverage breadth) at least in one sample in the cellular and virion metagenomes, respectively (Fig. S6). We next explored the spatiotemporal distribution of CL500-11 viruses by metagenomic read mapping. We selected two high-coverage viral contigs (LB2301E_00302_ctg_1 and LB2301H_00224_ctg_1) (Fig. 2D) and a viral contig detected in 20 CL500-11 SAGs (LB2301E_00219_ctg_540) (Fig. 2C). Notably, the two high-coverage contigs were both circularly assembled and shared synteny (Fig. 5). By querying these contigs against reference viral contigs derived from metagenomes, we identified four viruses (LBV_5008, LBV_7831, LBVC_2167, and LBVC_669) that shared synteny to the two circular viruses (Fig. S7). The seven viruses and a long-read assembled circular MAG of the host (CL500-11) recovered previously from the lake (7) were together used as a reference for metagenomic read mapping. LB2301E_00302_ctg_1 and LBV_7831 showed higher read coverage than other viral contigs in the hypolimnion and during the mixing period in both the cellular and virion fractions (Fig. 6). LB2301E_00219_ctg_540 was subsequently abundant but showed disproportionally higher coverage in the cellular fraction than in the virion fraction (Fig. 6).

**Fig. 5.**
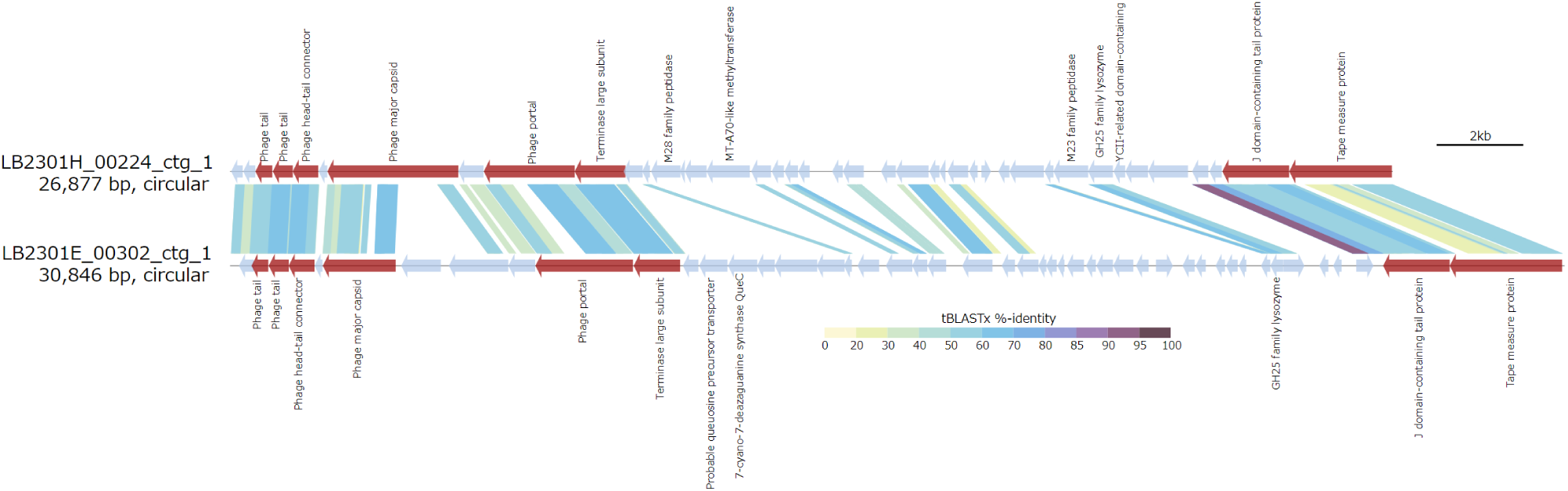
tBLASTx alignment of two circular CL500-11 viruses discovered in this study, LB2301H_00224_ctg_1 and LB2301E_00302_ctg_1. Arrows indicate predicted genes; labels indicate functional annotations; red indicates viral structural proteins. Sequences may be inversed or circularly permuted to show alignment more clearly.

**Fig. 6.**
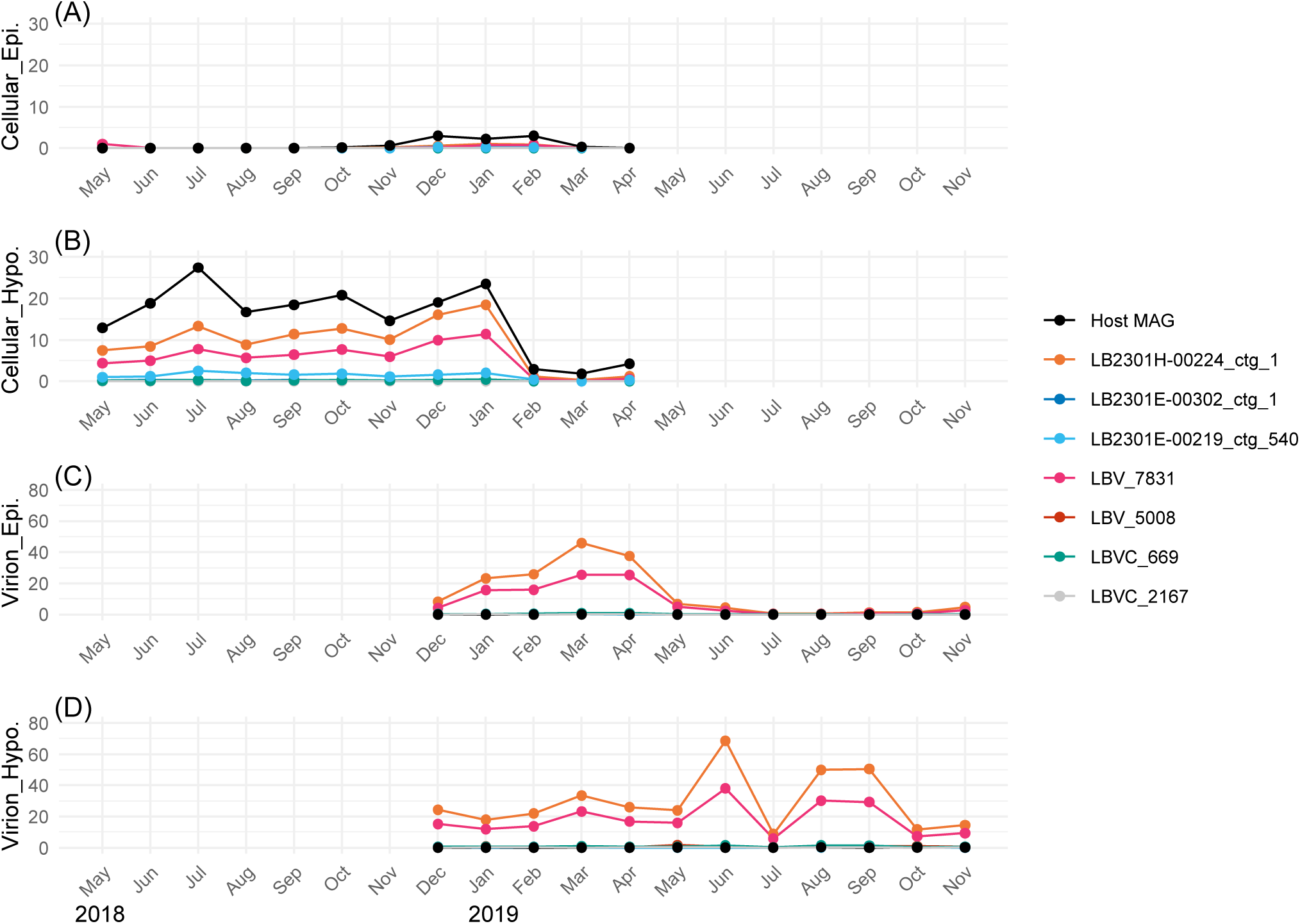
Monthly abundance dynamics of CL500-11 bacteria and their viruses (three contigs featured in the main text and their relatives in the database; Fig. S7) estimated by metagenomic read coverage. Coverages among the samples were normalized by mapping one million read pairs downsampled from each sample. Abundances in the (A, B) cellular fraction from May 2018 to April 2019 and (C, D) virion fraction from December 2018 to November 2019 in the epilimnion and hypolimnion, respectively. For the reference host genome, we used the circular CL500-11 long-read assembled metagenome-assembled genomes (MAG) generated in Lake Biwa (Okazaki et al., 2022).

## DISCUSSION

### SAG-gel recovered high-quality SAGs representing the lake bacterial community

Although limited assembly quality is a major problem in single-cell genomics (37–39), our SAGs generated high-quality assemblies (Fig. 1A and B), including a circularly assembled genome. These high-quality assemblies were attributable to the use of SAG-gel technology. In this technique, DNA extraction and whole-genome amplification of individual cells are performed within a gel bead, and only gel beads showing sufficient DNA amplification are sorted and sequenced, thus enabling efficient and high-quality reconstruction of SAGs with a reduced risk of contamination (17, 40). Their high quality is also attributable to the dominance of low-GC bacteria (e.g., Pelagibacteraceae and Nanopelagicaceae), facilitating less biased whole-genome amplification (41, 42). Indeed, our high-quality SAGs were enriched in low-GC genomes (Fig. 1A). The phylogenetic composition of the SAGs was generally in agreement with the known composition of the bacterial community in the lake; members of Pelagibacteraceae and Nanopelagicaceae generally dominate across depths and seasons, whereas CL500-11 shows a preference for the hypolimnion and is absent in the summer epilimnion (7, 9). The occurrence of CL500-11 in the winter epilimnion can be explained by the dispersal of cells by the onset of winter vertical mixing (43). The other dominant members (Ilumatobacteraceae, Burkholderiaceae, Beijerinckiaceae, Chitinophagaceae, and Methylophilaceae) are also abundant bacterioplankton lineages in Lake Biwa and other freshwater lakes (7, 44–46). Overall, our high-quality SAGs represented the diverse bacterioplankton community in the lake, allowing detailed single-cell-resolution investigation within and between lineages.

### Viral signals in SAGs revealed high viral diversity and novel virus–host pairs

Although we identified 176 viral contigs including 10 complete genomes from the SAGs, most were fragmented (Table S2), implying the presence of unassembled viruses. For more comprehensive viral detection by read mapping analysis, we used viral contigs from published Lake Biwa metagenomes. However, the SAG- and metagenome-derived viral contigs showed little overlap at the 95% ANI clustering threshold, and almost all metagenome-derived contigs were not detected by SAG read mapping (Fig. 2B), indicating their absence in the sequenced cells. Among the exclusively SAG-derived viruses, 73% and 24% were detected by cellular and virion metagenomic read mapping, respectively (Fig. S6). Their metagenomic detection indicates their presence in the samples, although they were unassembled in the metagenomes, presumably due to their low abundance or high genomic microdiversity, which hinders metagenomic assembly (47–49). By contrast, SAG-derived viruses not detected by metagenomic read mapping indicate their absence in the metagenomic samples, possibly because of the temporal shift of the viral community in the lake, as the metagenomes were collected 3–4 years prior to the collection of our SAGs. Collectively, the gap between the SAG and metagenomic viromes indicated the viral community in the lake to be highly diverse.

An important achievement of this study is the identification of novel virus–host pairs by the detection of viral signals in SAGs, which is unprecedented in a freshwater lake. Although large-scale viral metagenomic surveys in freshwater lakes routinely recover thousands of viral genomes (6, 8, 10, 50), the hosts of most of these viruses are unknown, which limits understanding of their ecology. Furthermore, host prediction relies mainly on gene phylogeny, which does not resolve the host to a low taxonomic level. For example, the actinobacterial *whiB* gene contributes a significant portion of the host prediction in freshwater viromes (8, 10, 51), but it predicts the host only to the phylum level, which limits ecological interpretations given the broad diversity of *Actinobacteria* in freshwater ecosystems (11, 45, 52). Our results linked viruses and hosts at the species level (Fig. S5 and Table S2), identifying ecologically important viral species infecting ubiquitous and abundant bacterioplankton lineages, as demonstrated by the discovery of CL500-11 virus discussed later.

One could argue that detecting a virus in a SAG does not necessarily indicate a specific interaction and can result from non-specific attachment of a virus to a cell or random co-encapsulation of a free-living virion and cell during single-cell isolation. In fact, some of our predictions were validated by known virus–host relationships. For example, two viral contigs assembled in the Pelagibacteraceae SAGs (LB2206E_00171_ctg_26 and LB2301E-00765_ctg_133) were relatives of HTVC010P (Fig. S8), a known Pelagibacteraceae virus in marine and freshwater systems (53, 54). Moreover, most of the viruses detected in multiple SAGs were consistently detected in the same genus (Fig. 2C), precluding random associations. However, in a few cases, the same virus was detected in multiple genera of SAGs (Fig. 2C), and an apparent false positive occurred, in which fragments of viral genomes closely related to known T4-like cyanophage S-SM2 (55) were detected in a Nanopelagicaceae SAG (Figs. 4C and S9). In conclusion, although the detection of a virus in a SAG generally reflects a *bona fide* virus–host association, the consistency of the data should be considered because of the possibility of false positives.

We characterized high-coverage viral contigs to differentiate viruses within a lytic cycle (i.e., high copy number in a cell). However, as reported previously (15), the assembly coverage of the contigs in an SAG exhibited broad and continuous distribution (Fig. 2D) due to the bias introduced by whole-genome amplification (41, 42). Thus, the assembly coverage did not enable objective definition of active viruses, except for the two CL500-11 circular viruses, as discussed below.

### Infection rate of the lake bacterioplankton community

The proportions of SAGs with a virus (12.1–18.1%; Fig 3A) were higher than those of visibly infected cells (0.9–4.1%) previously reported by transmission electron microscopy in Lake Biwa (56, 57). The higher rate obtained in this study suggests the high sensitivity of our read mapping-based approach to detect viral associations without virion formation. Nevertheless, our viral detection rates were lower than those in prior single-cell genomics studies (26% to >60%) (12, 13, 15, 16). This gap is attributable to differences in the diversity of the host community, because these studies targeted low-diversity communities or small numbers of taxa. It is also possible that the viral detection rate was underestimated because of the conservative threshold used in the viral identification pipeline. However, relaxing the viral identification criteria would introduce more mobile genetic element (MGE)-like sequences due to the presence of virus-like genes in some MGEs, as is the case of LB2301E_00219_ctg_540 discussed below. Therefore, leveraging the high-quality assembly of our SAGs, we applied conservative thresholds by using the consensus of three virus identification tools to limit our analysis to confidently identified dsDNA phages. Notably, the viral detection rate of Pelagibacteraceae (7.1%; Fig. 3B) was comparable to those determined for marine Pelagibacteraceae and their abundant phages by fluorescence *in situ* hybridization (58).

Co-infection of the same host cell by multiple viruses facilitates cooperation, competition, and co-evolution among viruses (59, 60). In this study, 55 SAGs detected multiple viral contigs (Fig. 3A and Table S3). However, only one SAG detected contigs that likely originated from multiple viral genomes (Fig. S2), and the other cases resulted from fragmented assemblies of the same virus (Fig. S3) or were not characterized due to inadequate assembly length and the lack of marker genes (Table S3). Thus, evaluation of the frequency and ecological implications of co-infection warrants a larger-scale study that compensates for the low viral detection rate.

### Contrasting viral defense strategies of oligotrophs and copiotrophs

The viral infection rate was stable across four samples from different seasons and water layers (Fig. 3A), but showed greater variance among host lineages (Fig. 3B), suggesting that susceptibility to viral infection depends on the host lineage. Notably, copiotrophic bacterial lineages had higher viral detection rates than oligotrophic lineages. Oligotrophs (i.e., K-strategists) are characterized by slow growth and persistent dominance in pelagic open water systems by adapting to stable and resource-poor conditions. By contrast, copiotrophs (i.e., r-strategists) are characterized by rapid growth and opportunistic dominance under resource-rich conditions and have a versatile metabolism, enabling their survival under competitive conditions (44, 61, 62). Specifically, members of the Pelagibacteraceae (also known as LD12 or SAR11), Nanopelagicaceae (acI), Ilumatobacteraceae (acIV), and Methylophilaceae (LD28), which are typical oligotrophs in freshwater ecosystems (44, 63), showed lower viral detection rates than the Burkholderiaceae and Chitinophagaceae (Fig. 3B), typical copiotrophs dominating under resource-rich conditions such as algal blooms (44, 63). The highest viral detection rate was for Beijerinckiaceae, most of which were affiliated with a species of *Methylocystis* (Figs. 3B and S5), which are aerobic methane-oxidizing bacteria (64). They have been regarded to be copiotrophs in the lake based on their non-persistent occurrence (7, 9), likely reflecting the highly spatially and temporally heterogeneous methane concentration in the lake (65).

Intriguingly, the high viral detection rates of copiotrophs resulted not from the domination of a single virus but from collective infection by diverse viruses (Fig. 4A, B). Infection by diverse viruses suggests weak density-dependent virus–host selection, because a lower infection rate (resistant hosts are selected) or occupancy of a few viruses (viruses breaking host resistance are selected) is expected under strong density-dependent virus–host selection. We argue that the growth of copiotrophs is primarily controlled by resource availability, making their interaction with viruses non-persistent. Thus, the host population collapses due to resource deficiency before density-dependent virus–host selection occurs. The strategy of the host is to invest more in resource competition to outgrow rather than defend against viruses (Fig. 7). This strategy is analogous to the greater vulnerability of freshwater copiotrophs to protistan grazing because they have larger cells and compensate for grazing loss with a higher growth rate than that of oligotrophs (63, 66). Overall, our results support the notion that copiotrophs are primarily coping with bottom-up factors (competition for resources), and that fluctuations in their abundance prevent selective interactions with top-down factors (viral lysis and protistan grazing).

**Fig. 7.**
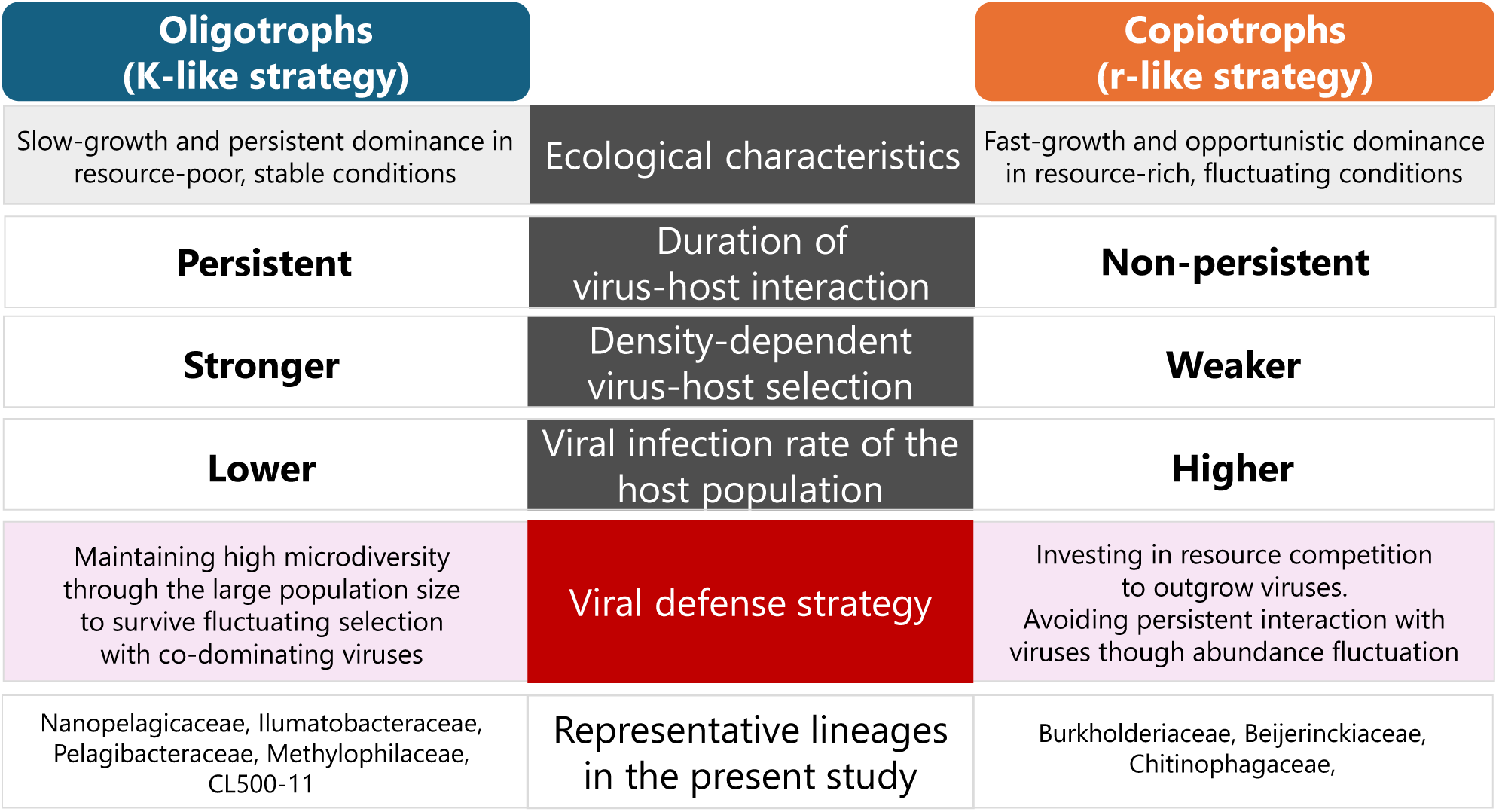
Contrasting viral defense strategies of oligotrophs and copiotrophs proposed based on the results. Oligotrophs cope with stronger density-dependent virus–host selection due to persistent co-dominance with their viruses. Genome microdiversification to diversify virus–host specificity is an effective strategy for oligotrophs due to their large population size, which explains the low infection rate of their population. By contrast, copiotrophs undergo weaker density-dependent virus–host selection via non-persistent interactions with viruses due to their fluctuating abundance depending on resource availability. Therefore, the strategy of copiotrophs is to invest more in resource competition to outgrow viruses, rather than coping with the virus–host arms race, which explains the high rates of infection of copiotrophs by diverse viruses.

Oligotrophs exhibited a low viral detection rate (Fig. 3B) despite their persistence, indicating resistance to viral infection. Streamlined genomes of oligotrophs harbor fewer genetic defense systems such as CRISPR/Cas (11, 61, 62), and their viral defense is thought to be achieved by a high intra-species microdiversity (2, 5, 11). For example, hypervariable region 2 (HVR2) of the Pelagibacteraceae is enriched for glycosyltransferases and is thought to be responsible for generating the diversity of cell surface glyco-structures to evade phage recognition (67–70). Indeed, a pangenome graph of our Pelagibacteraceae SAGs demonstrated that their HVR2 regions were nearly unique to individual cells (Fig. S10), as also reported in a recent study of marine Pelagibacteraceae (71). Unlike the fluctuating abundance of copiotrophs, oligotrophs are less likely to be affected by population bottlenecks and thus maintain their genetic microdiversity (7). High microdiversity has been reported for viruses of oligotrophs, likely as a consequence of persistent fluctuating selection (72–75). Overall, our results highlight the contrasting viral defense strategies of oligotrophs and copiotrophs, inviting further investigation of their interactions with viruses (Fig. 7).

### SAGs revealed dominant CL500-11 viruses and their infection dynamics

This study is the first to identify a virus infecting CL500-11 (phylum Chloroflexi), an uncultured bacterioplankton lineage dominant in the oxygenated hypolimnion of deep freshwater lakes globally (23, 24, 76). CL500-11 is persistently dominant in the hypolimnion during the stratification period (24, 43), hinting at its oligotrophic lifestyle. Besides, their abundance decreases with the collapse of the thermocline during winter mixing (24, 43), prompting the investigation of viruses as potential mortality factors. We detected 17 viral contigs from 25 CL500-11 SAGs, among which the same viral contig was detected in 20 SAGs, and 14 viral contigs were detected in three SAGs (Fig. 4D). The other two viral contigs were circularly assembled, shared synteny, and were detected in two different CL500-11 cells collected in winter (Figs. 4D and 5). Together with their relatives in the reference virome (Fig. S7), these viruses showed a spatiotemporal distribution following that of the host in the metagenomes (Fig. 6), suggesting them to be *bona fide* CL500-11 viruses.

The two circular CL500-11 viruses exhibited far the highest assembly coverage than other chromosomal contigs in the SAG (Fig. 2D). Such an extreme viral contig coverage in a SAG was not observed in a prior study (15) and suggests lytic activity and a high burst size. Moreover, LB2301H_00224_ctg_1, one of the circular viruses, showed high metagenomic read coverages in the cellular and virion fractions in the hypolimnion throughout the water stratification period (Fig. 6), indicating their active replication in a cell and release of virions to the water column. Intriguingly, their metagenomic read coverage in the cellular fraction was close to that of their host in the hypolimnion at the end of stratification period (Fig. 6), although the virus was detected only in one SAG among the 215 CL500-11 SAGs analyzed (Fig. 4D). Collectively, our results suggest that LB2301H_00224_ctg_1 is an abundant and active lytic virus with a high burst size that infects a minor fraction (< 1%) of the host population. Of note, previous studies in Lake Biwa observed CL500-11-like (large, curved) cells with full of virions inside using transmission electron microscopy (56, 57). As reported in other oligotrophs, genomic microdiversity may underly virus–host specificity in this system. Indeed, the microdiversity of CL500-11 was comparable (∼40,000 SNVs/Mb) to those of other oligotrophs according to metagenomic read mapping (7). In summary, the combination of SAG and metagenomic data revealed one of the most quantitatively significant virus–host pairs discovered in freshwater systems and the marked heterogeneity of their interactions, which has been overlooked in metagenomics studies.

Another CL500-11 viral contig, LB2301E_00219_ctg_540, was the most prevalent viral contig in this study, being detected in 20 SAGs (Figs. 2C and 4D). The contig showed detectable metagenomic read coverage in the cellular fraction, but was absent from the virion fraction (Fig. 6). It encoded an integrase and showed homology to a portion (inserted between tRNAs) of a CL500-11 MAG assembled in a previous study, in which complex structural variants were identified by long-read metagenomic read mapping (7) (Fig. S11). LB2301E_00219_ctg_540 is dominated by repetitive proteins and harbors only one viral marker gene (a phage head morphogenesis protein) in nested structural variants (Fig. S11). Collectively, we concluded that LB2301E_00219_ctg_540 is an MGE identified as a viral contig based on the presence of an auxiliary viral structural-like protein. Although MGE is beyond the scope of this study, the dissemination of the integrated MGE within the persistent and microdiverse host population is intriguing. We propose that the function of the enriched genetic structural variant (Fig. S11) is the key to understanding the mechanism underlying its persistent co-dominance with its host.

### Conclusion

We discovered diverse and heterogeneous virus–host interactions, which have been overlooked in metagenomics studies. The contrasting viral defense strategies of oligotrophs and copiotrophs (Fig. 7) provided a framework for further studies of these virus–host interactions. We propose that the persistence of density-dependent interactions is the key to understanding the diverse viral defense strategies and mechanisms underlying virus–host co-existence. The facts that our SAGs represented a minor portion of the metagenomic viral diversity (Fig. 2B), that most viruses were detected in a single SAG (Fig. 2C), and that only a single putative co-infection was identified (Fig. S2, Table S3), together indicate that our understanding of the virus–host network in Lake Biwa is inadequate. Therefore, further research should sequence greater numbers of cells and samples or focus on particular lineages using, for example, a targeted single-cell genomics approach (77). Extending the research beyond metagenomic resolution will advance our understanding of virus–host interactions, which are among the most important drivers of microbial biogeochemical cycles and genomic diversification.

## Data availability

The raw sequencing reads for the 862 SAGs are available under BioProject ID PRJDB18380. The nucleotide fasta files of the assembled 862 SAGs are available at https://doi.org/10.6084/m9.figshare.26243444

## Acknowledgments

We thank Shang Shen for providing the LBV dataset from their published work. This work was supported by Center for Ecological Research, Kyoto University, a Joint Usage/Research Center, MEXT KAKENHI (22H00382, 22K15182, 23K13612), JST FOREST (JPMJFR2273), and JST ACT-X (JPMJAX20BE). Computation time was provided by the SuperComputer System, Institute for Chemical Research, Kyoto University.

---

**Fig. S1.**
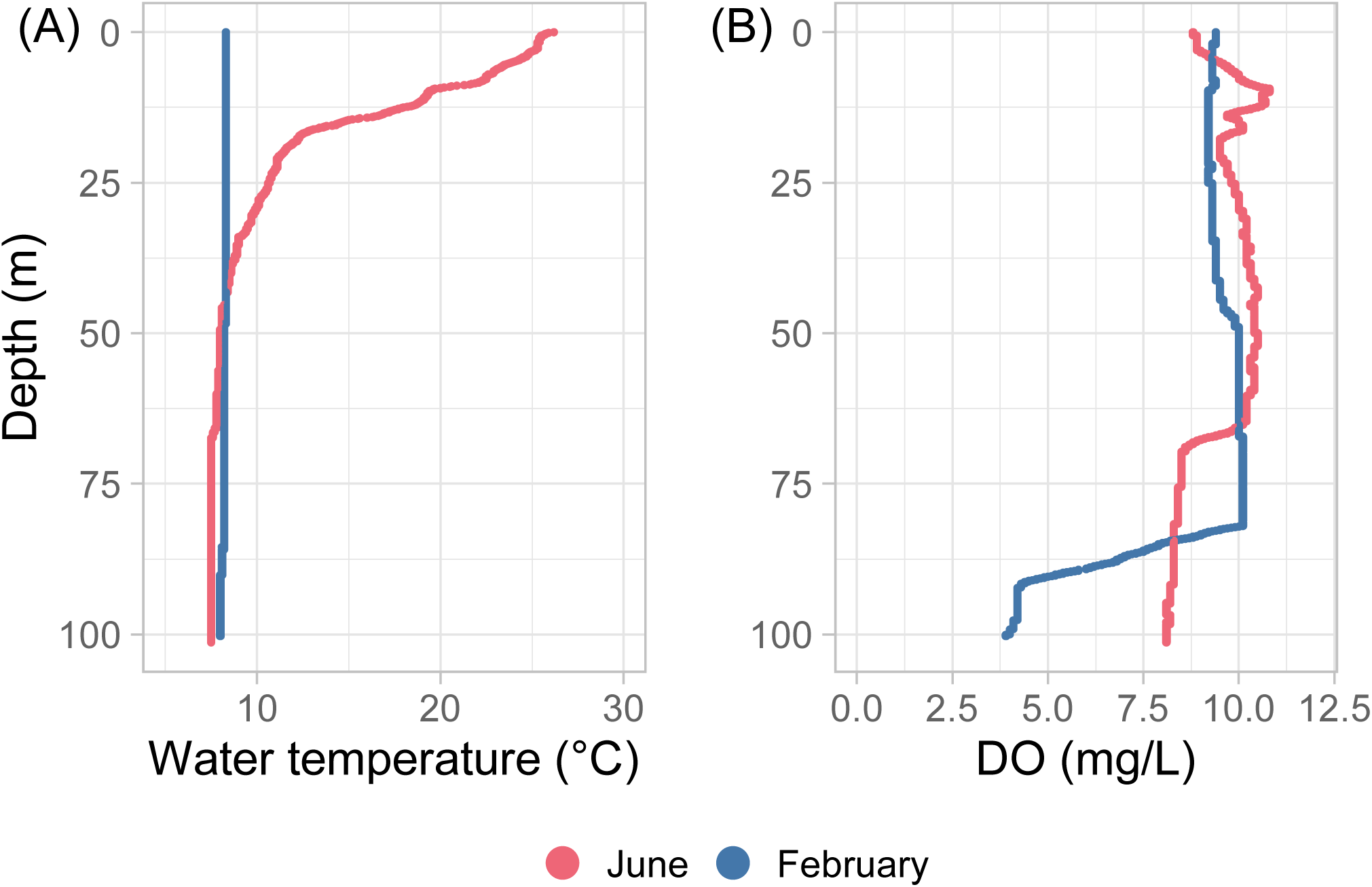
Vertical profiles of (A) water temperature and (B) dissolved oxygen (DO) concentration at the time of sampling (June 29, 2022 and February 1, 2023).

**Fig S2.**
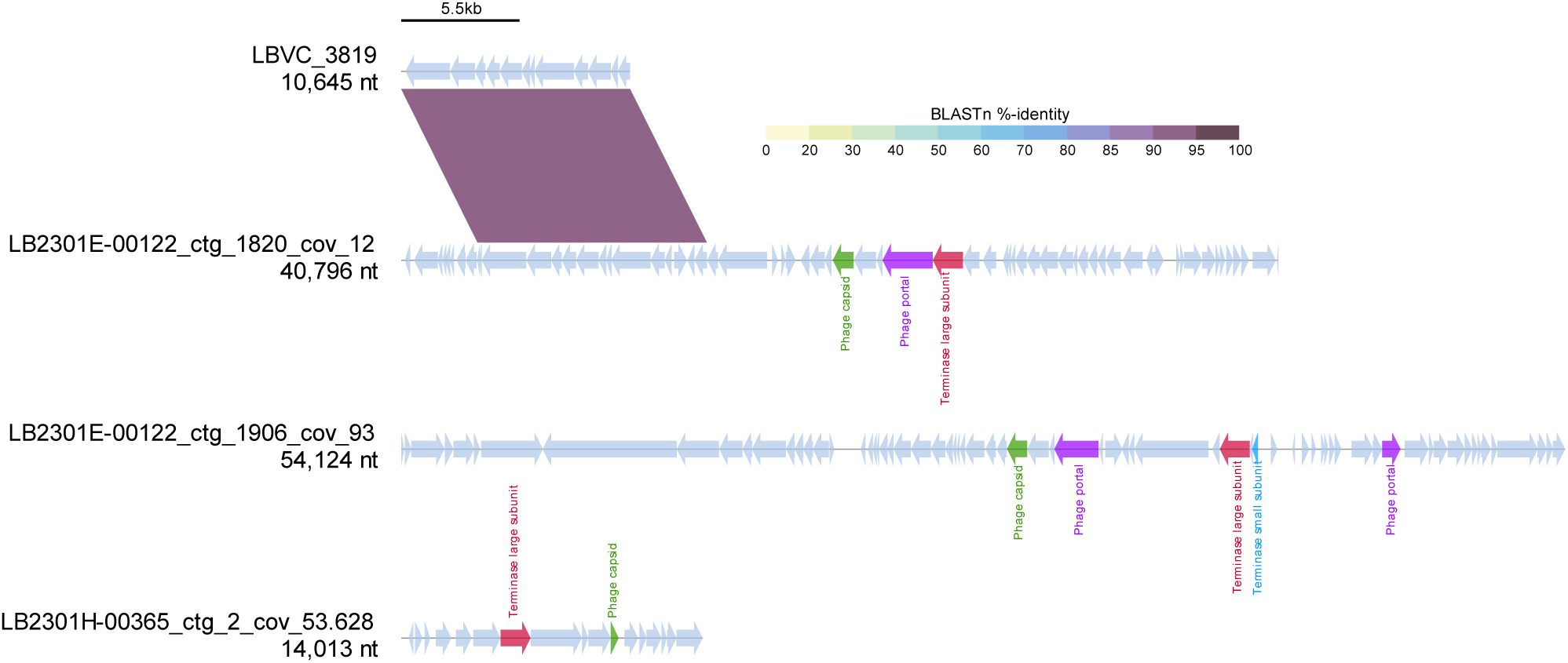
BLASTn-alignment of four viral contigs detected in the same SAG (LB2301E-00122). Arrows indicate predicted genes; colors indicate viral structural genes. Three viral contigs with overlapping structural proteins showed no homology, indicating the detection of multiple viral genomes from a single cell. The sequences may be inversed to show alignment more clearly.

**Fig. S3.**
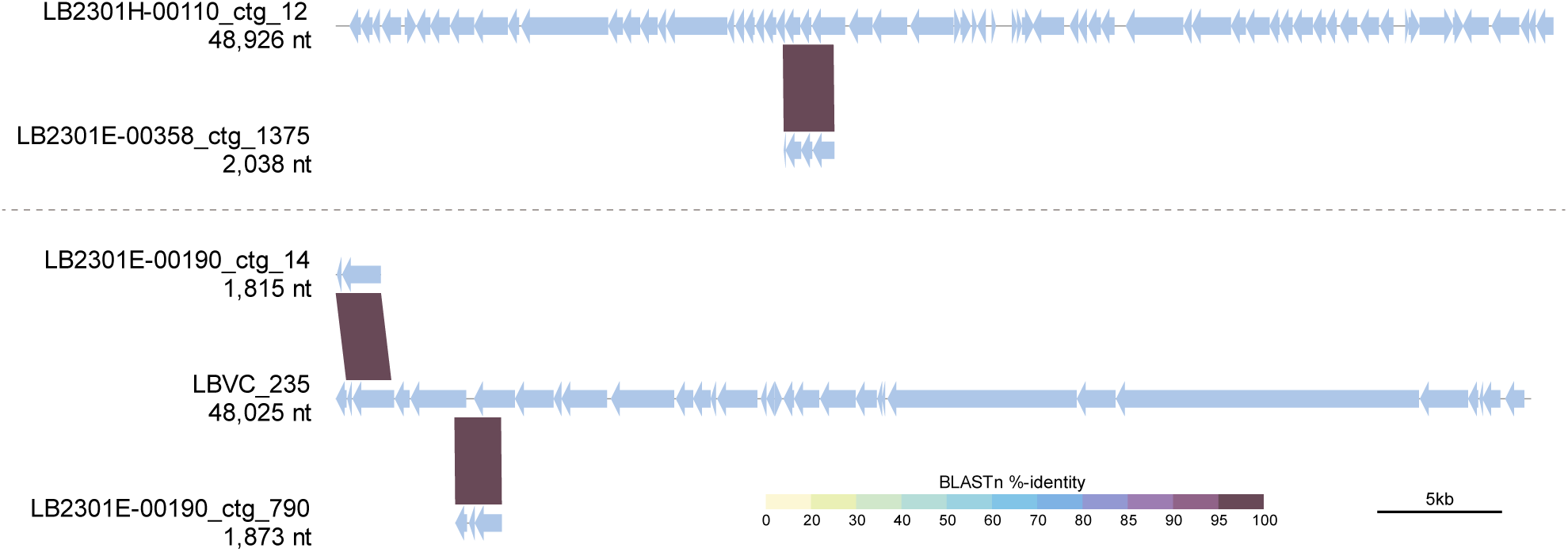
BLASTn alignment of viral contigs co-detected in two SAGs, LB2301H-00110 (top two sequences) and LB2301E-00190 (bottom three sequences). Short contigs are aligned to a much longer contig, suggesting that they originated from the same viral genome but eluded de-replication because the aligned fraction size was lower than the threshold (10%) set in dRep software. Sequences may be inversed to show alignment more clearly.

**Fig. S4.**
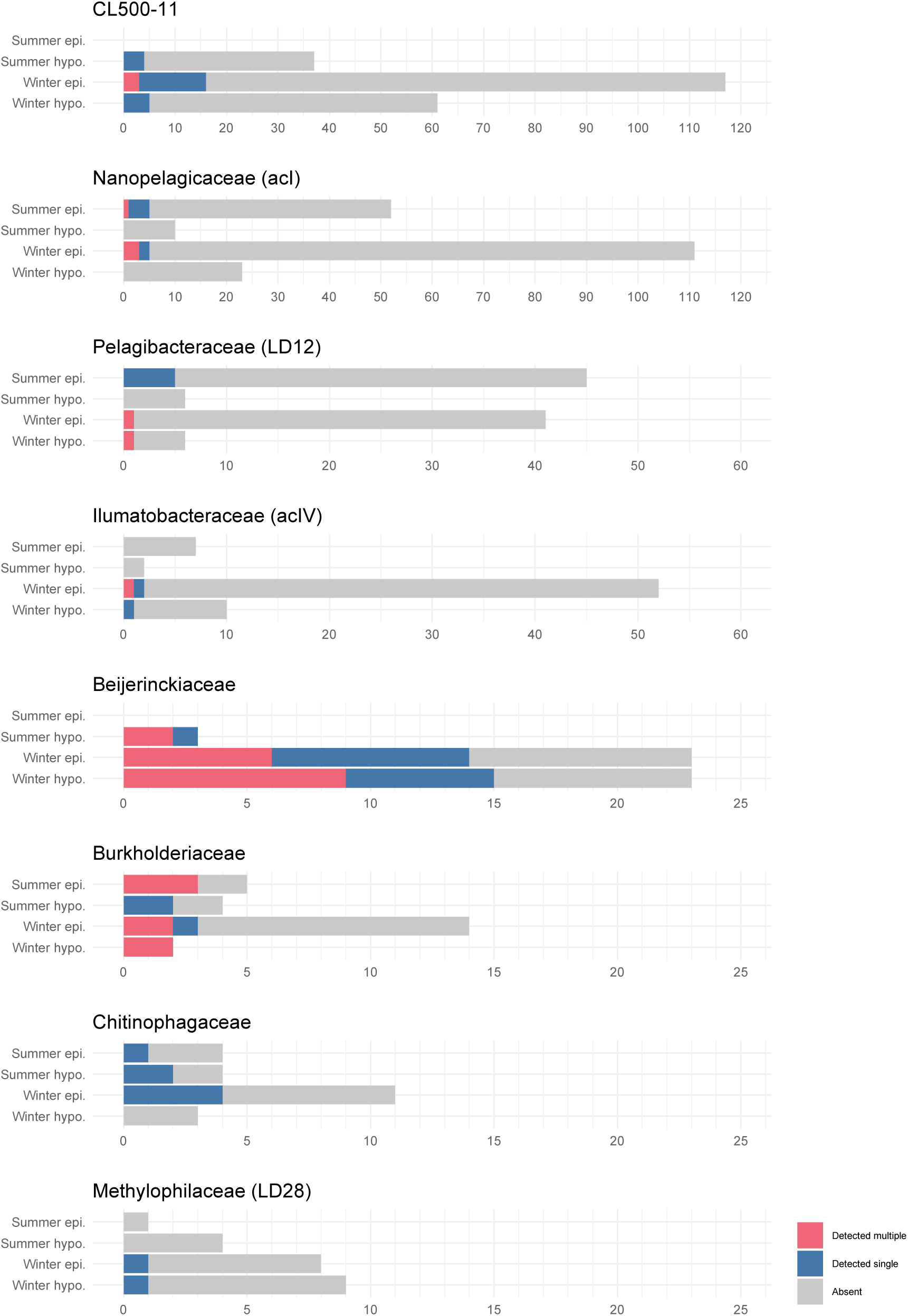
Numbers of SAGs with viral detection for each sample in each family. The detection of single or multiple viral contigs is indicated by different colors.

**Fig. S5.**
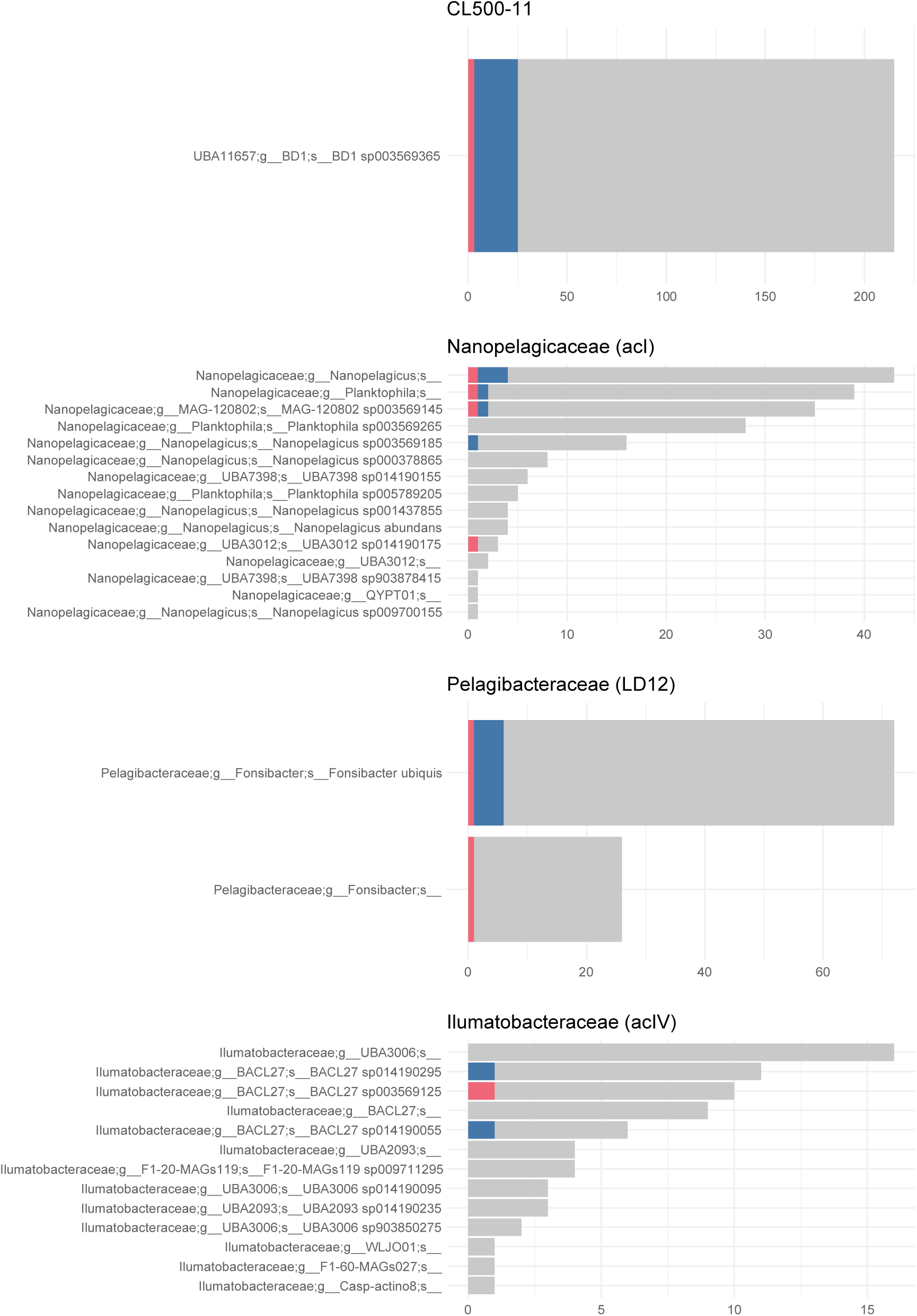
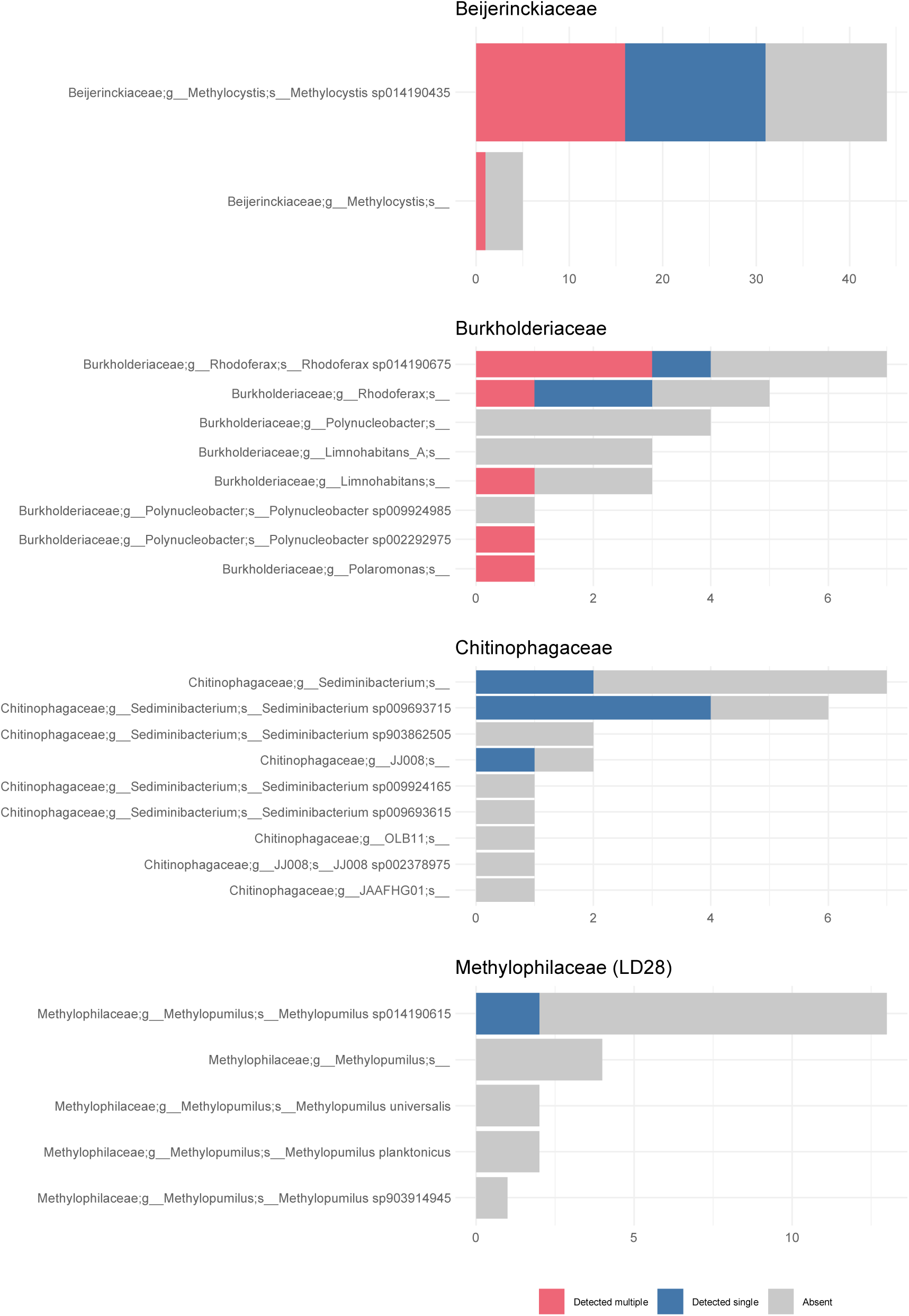
Numbers of SAGs with viral detection for each species in each family. The detection of single or multiple viral contigs is indicated by different colors.

**Fig S6.**
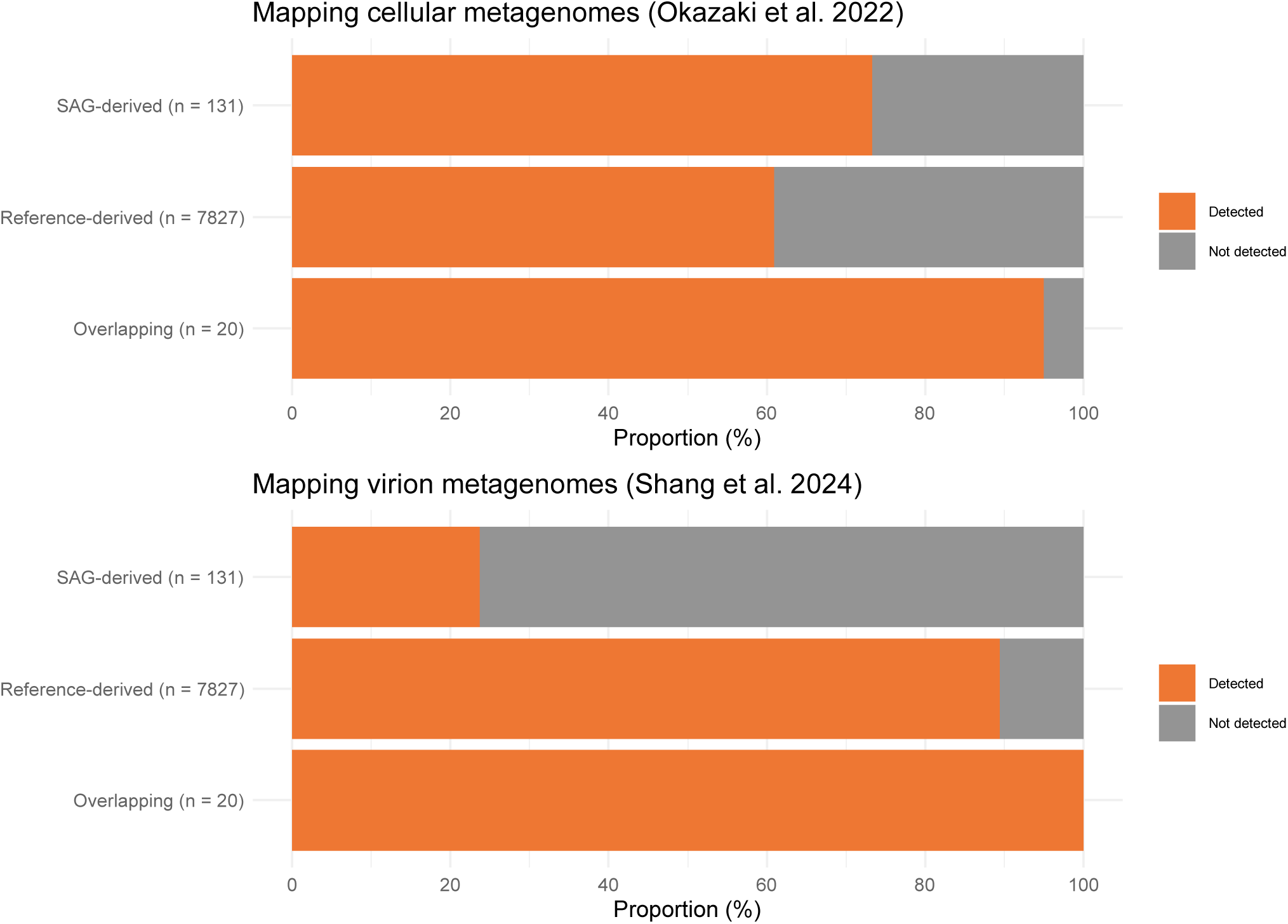
Proportions of viruses detected by metagenomic read mapping in cellular and virion metagenomes. If one of the 24 metagenomic samples in each size fraction showed a > 50% mapping breadth in a contig (i.e., > 50% of a contig was covered by mapped reads), the virus was regarded as detected in the size fraction. Viruses are grouped into three categories, as in Fig. 2B.

**Fig. S7.**
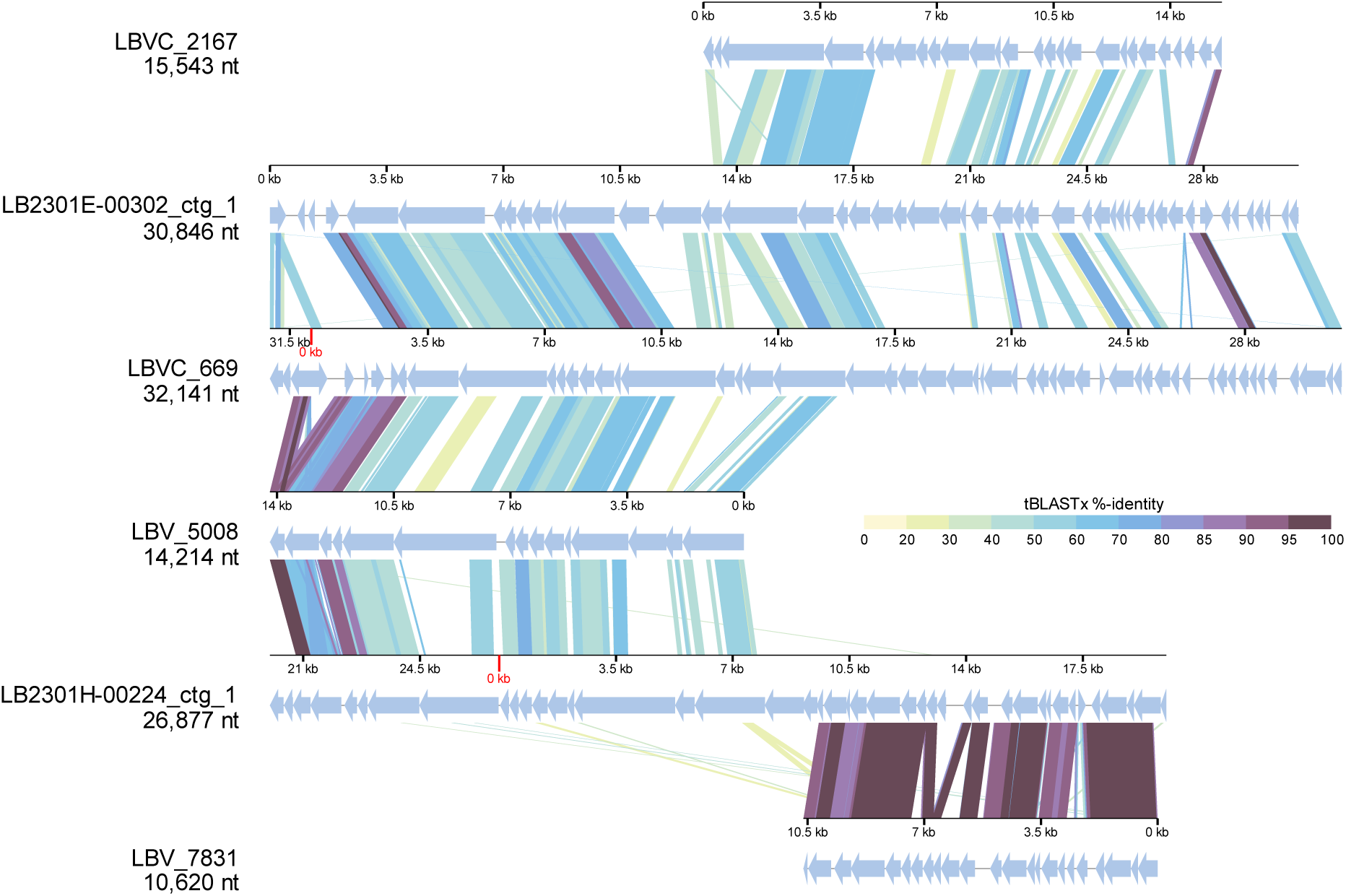
tBLASTx alignment of the two circular CL500-11 viruses (LB2301H_00224_ctg_1 and LB2301E_00302_ctg_1) and their related reference viruses. Sequences may be inversed or circularly permuted to show alignment more clearly.

**Fig. S8.**
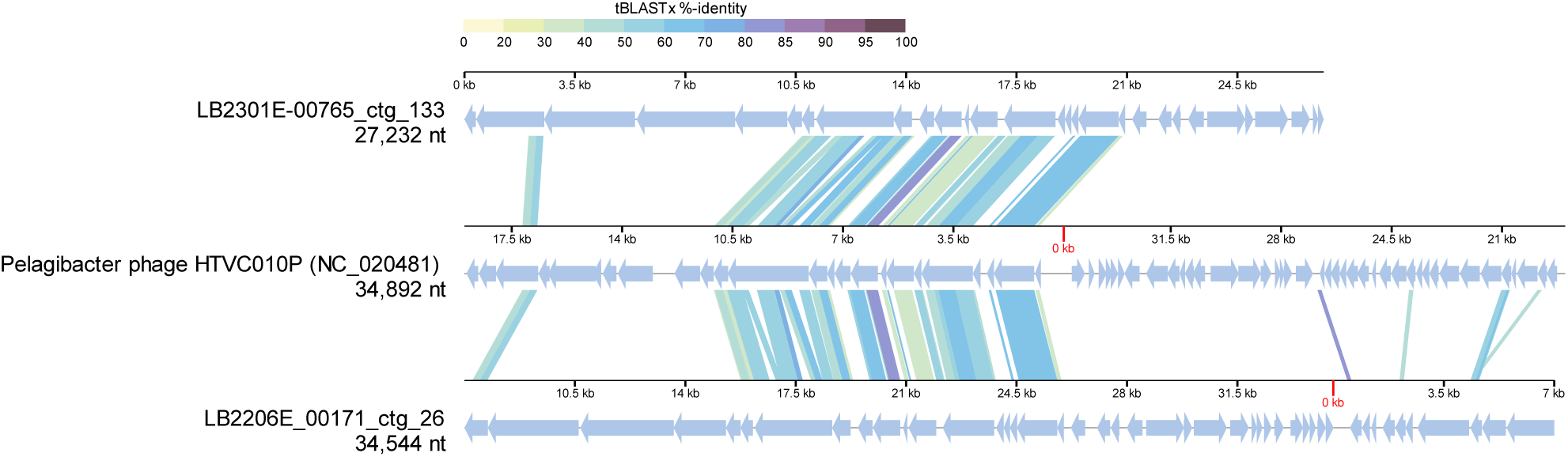
tBLASTx alignment of the Pelagibacter phage HTVC010P and its related viral contigs (LB2301E_00765_ctg_133 and LB2206E_00171_ctg_26) recovered from Pelagibacteraceae SAGs. Sequences may be inversed or circularly permuted to show alignment more clearly.

**Fig. S9.**
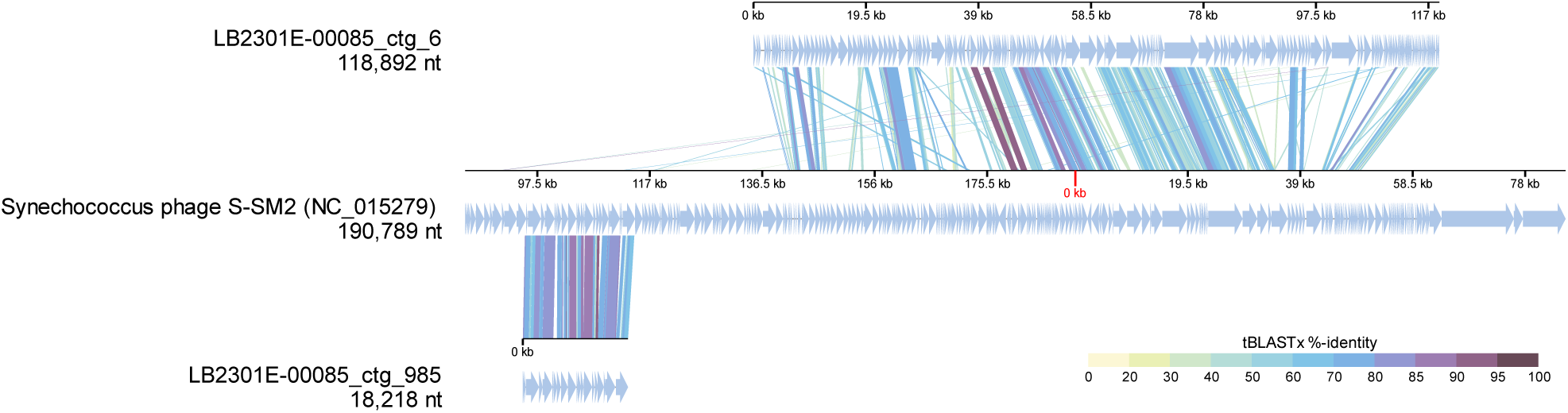
tBLASTx alignment of *Synechococcus* phage S-SM2 and its related viral contigs recovered from LB2301E-00085, an actinobacterial SAG. Sequences may be inversed or circularly permuted to show alignment more clearly.

**Fig. S10.**
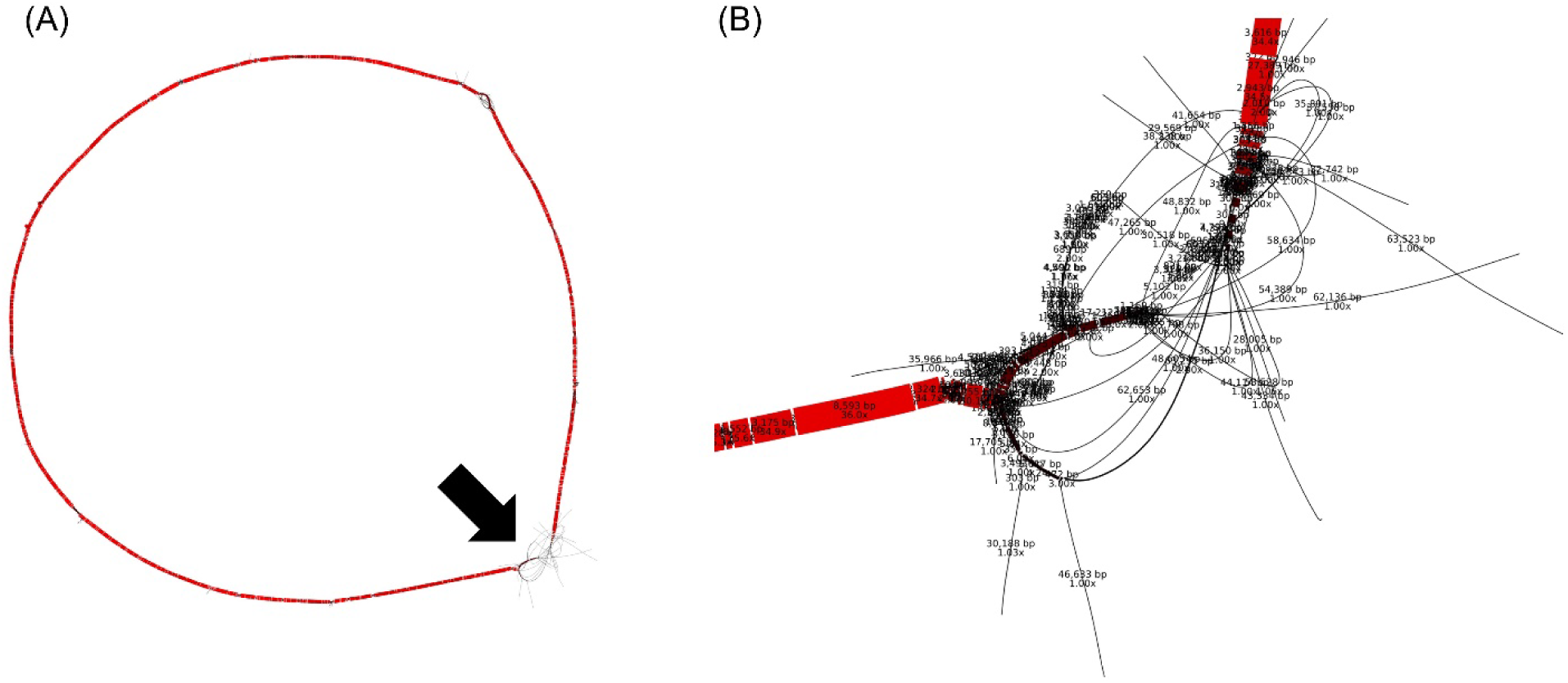
Pangenome graph of 39 high-quality (QS > 50) SAGs affiliated with *Fonsibacter ubiquis* collected from the summer epilimnion. Graph was generated by SuperPang using the default parameters and visualized by Bandage; genomic segments sharing > 95% similarity are shown as non-branching paths. Thus, inconsistent sequences between SAGs with < 95% sequence similarity are shown as branches in the graph. The width of a path indicates its coverage (average prevalence of its constituent kmers in input genomes). (A) Structure of the pangenome graph. Arrow indicates hypervariable region 2 (HVR2). (B) Enlarged HVR2 in the pangenome graph. Path labels indicate length and coverage.

**Fig. S11.**
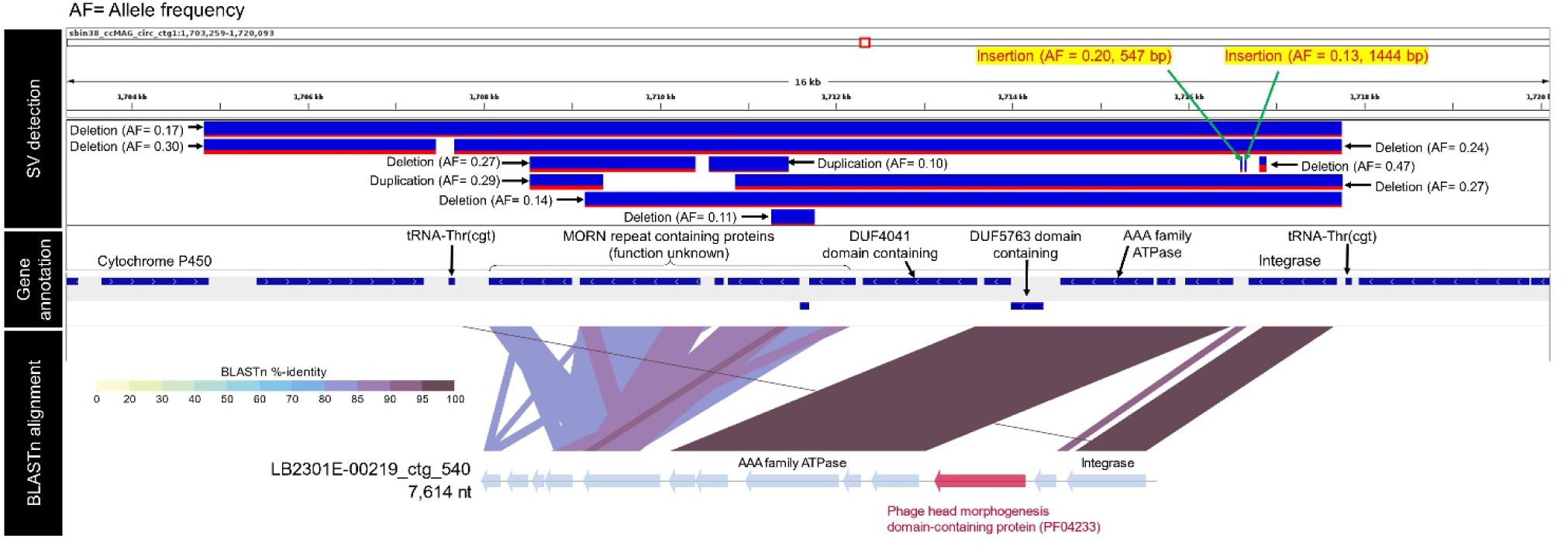
The contig LB2301E-00219_ctg_540 aligned to a portion of a MAG of the host (CL500-11) shown with structural variants detected by metagenomic long-read mapping (Okazaki et al. 2022). The host MAG was assembled into a single circular contig; the segment from 1703 to 1720 kb is shown. The top layer shows the positions of detected structural variants (deletions, insertions, and duplications) with estimated allele frequencies. Insertions are shown with their estimated lengths. The middle layer shows gene annotations of the MAG in the original study (Okazaki et al. 2022). The bottom layer shows the BLASTn alignment between LB2301E-00219_ctg_540 and the MAG. Arrows indicate predicted genes. Red arrow indicates a phage head morphogenesis protein, which is absent in the MAG but present in the predicted insertion.

## Legends for supplementary tables

Table S1. Statistics of 1657 assemblies with a > 1-kb contig generated from individual gel beads. In total, 862 medium- or high-quality single-cell amplified genomes (SAGs) were obtained and used in downstream analyses.

Table S2. Statistics of 176 viral contigs assembled from 85 SAGs, and detected as dsDNA phages by at least two of geNomad, VIBRANT, and VirSorter2 (see main text for detection criteria for each tool).

Table S3. Statistics of co-detected viral contigs. Two to nine viral contigs were co-detected in each of 55 SAGs, as indicated by different colors in the first and second columns of the table.

